# Frataxin deficiency disrupts mitochondrial respiration and pulmonary endothelial cell function

**DOI:** 10.1101/2022.08.22.504849

**Authors:** Miranda K. Culley, Monica Mehta, Jingsi Zhao, Dror Perk, Yi Yin Tai, Ying Tang, Sruti Shiva, Marlene Rabinovitch, Mingxia Gu, Thomas Bertero, Stephen Y. Chan

## Abstract

Deficiency of iron-sulfur (Fe-S) clusters promotes metabolic rewiring of the endothelium and the development of pulmonary hypertension (PH) *in vivo*. Joining a growing number of Fe-S biogenesis proteins critical to pulmonary endothelial function, recent data highlighted that frataxin (FXN) reduction drives Fe-S-dependent genotoxic stress and senescence across multiple types of pulmonary vascular disease. Trinucleotide repeat mutations in the *FXN* gene cause Friedreich’s ataxia, a disease characterized by cardiomyopathy and neurodegeneration. These tissue-specific phenotypes have historically been attributed to mitochondrial reprogramming and oxidative stress. Whether FXN coordinates both nuclear and mitochondrial processes in the endothelium is unknown. Here, we aim to identify the mitochondria-specific effects of FXN deficiency in the endothelium that predispose to pulmonary hypertension. Our data highlight an Fe-S-driven metabolic shift separate from previously described replication stress whereby FXN knockdown diminished mitochondrial respiration and increased glycolysis and oxidative species production. In turn, FXN-deficient endothelial cells exhibited a vasoconstrictive phenotype consistent with PH. These data were observed in both primary pulmonary endothelial cells after pharmacologic inhibition of FXN and inducible pluripotent stem cell-derived endothelial cells from patients with FXN mutations. Altogether, this study defines FXN as a shared upstream driver of pathologic aberrations in both metabolism and genomic stability. Moreover, our study highlights FXN-specific vasoconstriction, suggesting available and future therapies may be beneficial and targeted for PH subtypes with FXN deficiency.

**Graphical Abstract:** 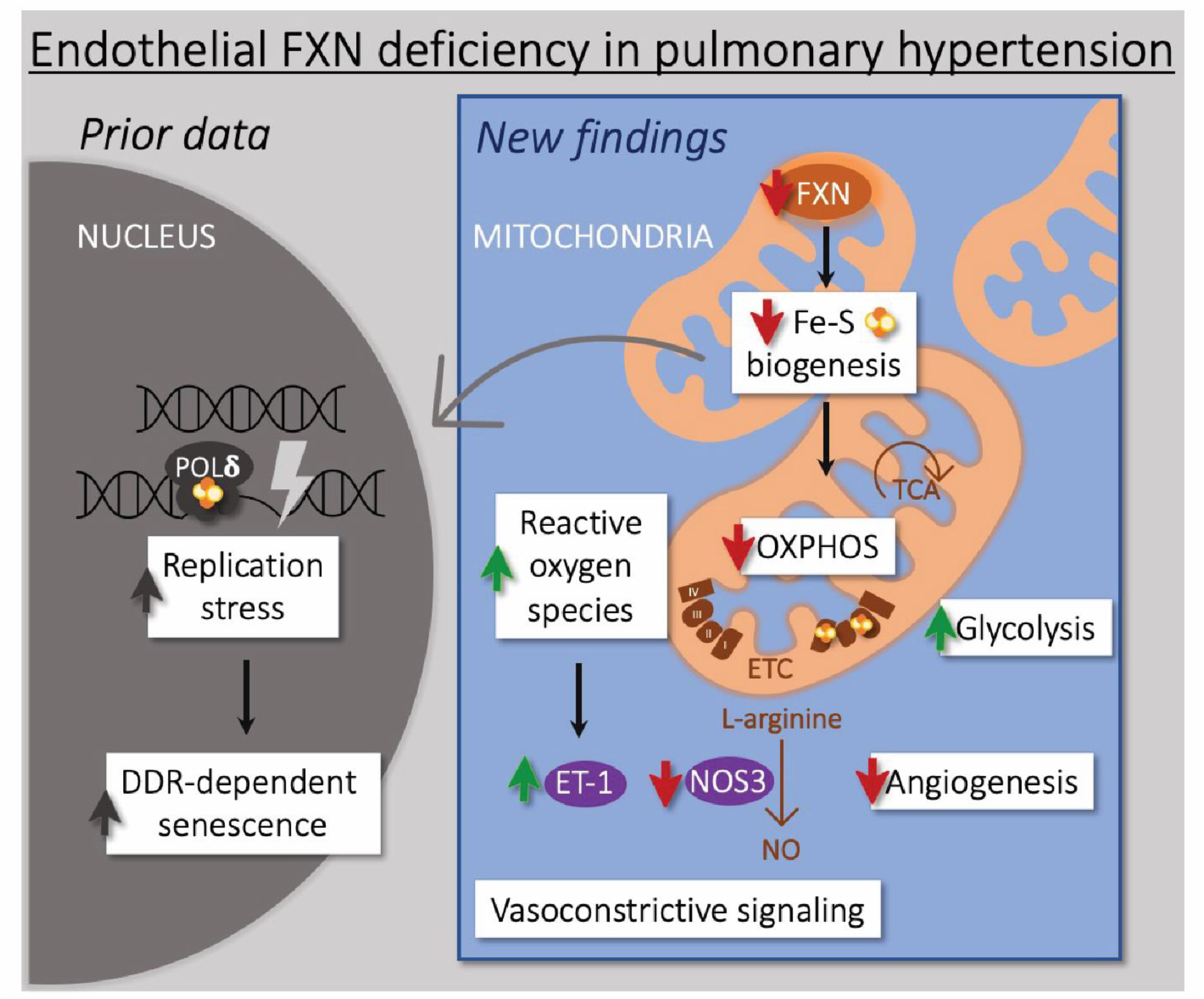

## Introduction

Pulmonary hypertension (PH) is a progressive and fatal disease of the lung vasculature in which mitochondrial dysfunction contributes to pathogenesis [1]. A hallmark of the metabolic reprogramming observed in the endothelium in PH is the shift from oxidative phosphorylation to glycolysis [2-5], similar to the Warburg effect in cancer [6, 7]. Iron-sulfur (Fe-S) cluster deficiency contributes to the disruption of endothelial cell metabolism and PH *in vivo* [8-10]. Fe-S clusters are essential, redox-capable cofactors incorporated into different apoproteins across cellular compartments [11]; for example, Fe-S centers facilitate electron transfer between mitochondrial complexes allowing for effective respiration [12]. Our previous data established that both acquired and genetic deficiencies of the Fe-S biogenesis genes *ISCU1/2* and *BOLA3* resulted in metabolic reprogramming in the form of the reduced respiration with a concomitant increase in aerobic glycolysis [8-10], reactive oxygen species (ROS) formation [8, 10], and lipoate-dependent regulation of glycine [10]. These Fe-S-specific metabolic changes promoted shared endothelial-specific phenotypes relevant to PH including increased vasoconstrictive signaling and dysregulated angiogenesis, ultimately resulting in pulmonary vascular disease *in vivo* [8-10].

Separately, CRISPR/Cas9 genome editing was used to introduce a point mutation in the Fe-S biogenesis gene *NFU1* (NFU1^G206C^) in Sprague-Dawley rats. The lungs from this model, and specifically pulmonary artery smooth muscle cells [13], exhibited decreased electron transport chain complex expression and activity and both histologic and hemodynamic changes consistent with PH [14]. How other Fe-S biogenesis gene deficiencies contribute to metabolic reprogramming and their dynamic consequences for endothelial cell phenotypes in PH has been incompletely studied.

We recently demonstrated that endothelial deficiency of frataxin (FXN), another nuclear-encoded, mitochondrial Fe-S biogenesis gene [15, 16], predisposed to PH development *in vivo* [17]. Specifically, FXN knockdown incited Fe-S-dependent replication stress and acute growth arrest that eventually precipitated endothelial senescence, which in turn, promoted vessel inflammation and remodeling consistent with pulmonary vascular disease. Of note, acquired FXN deficiency was linked with elevated senescence across models of Group 1, 2, and 3 PH with *in vitro* evidence that hypoxia-inducible factor-alpha (HIF-α) is responsible for transcript repression in hypoxic and inflammatory conditions. At the same time, inducible pluripotent stem cell-derived endothelial cells (iPSC-ECs) with biallelic *FXN* mutations exhibited the same mechanistic evolution toward senescence [17]. Coupled with rare lung samples from patients with genetic FXN deficiency, our data suggested PH may represent a previously underappreciated clinical phenotype in patients with Friedreich’s ataxia – a disease driven by trinucleotide repeat mutations in the *FXN* gene [18] and defined by severe neurodegeneration and cardiomyopathy [19].

While our study focused on the genotoxic effects of pulmonary endothelial FXN reduction, data in non-vascular cell types, specifically those with high bioenergetic capacity such as the nervous system, myocardium, and endocrine pancreas, have highlighted how FRDA-dependent FXN deficiency drives mitochondrial dysfunction [16]. Therefore, we investigated whether FXN deficiency controls Fe-S mediated metabolic reprogramming and endothelial dysfunction relevant to PH pathogenesis.

## Materials and Methods

### Cell culture

Primary human pulmonary artery endothelial cells (PAECs) and pulmonary artery smooth muscle cells (PASMCs) (PromoCell) were plated in collagen-coated plastic and cultured in cell-specific basal media supplemented with a growth media kit (PromoCell; Lonza) and 5% fetal bovine serum (FBS) without antibiotics or antifungals added. Experiments were performed between passages 4 to 11. Serum-starved primary cells were exposed to hypoxia (0.2% O_2_) at 37°C using a modular hypoxia chamber or standard non-hypoxic conditions (21% O_2_) at 37°C. Inducible pluripotent stem cells (iPSCs) from patients with FRDA (NIGMS Human Genetic Cell Repository at the Coriell Institute for Medical Research; GM23404, GM23913) and gender-matched controls (generously gifted by M. Gu and M. Rabinovitch) were differentiated into endothelial cells as previously described [20], grown in basal growth media (PromoCell) containing 20% FBS, and used for study between passages 2 and 8.

### Transfection

Transfection was performed in PAECs at 70-80% confluency using 40nM of FXN siRNA (Santa Cruz Biotechnology) or pooled negative control siRNA (Santa Cruz Biotechnology) and Lipofectamine 2000 reagent (Life Technologies) in one-part OptiMEM (ThermoFisher Scientific) and three-parts serum-starved cell-specific media (Lonza). Following 6-8 hours incubation, transfection media was removed and replaced with full-serum growth media. Experiments were completed 48 hours post-transfection.

### RNA extraction and quantitative PCR

Cells were lysed in Qiazol (Qiagen), and RNA extracted using Rneasy Mini Kit (Qiagen). Complementary DNA (cDNA) was synthesized via reverse transcription on an Applied Biosystems Real Time PCR instrument (ThermoFisher Scientific). Quantitative RT-PCR was performed on an Applied Biosystems QuantStudio 6 Flex Fast Real Time PCR device, and fold-change of RNA species was calculated using the formula (2^-ΔΔCt^) normalized to β-actin expression.

### Immunoblotting

Cells were lysed in RIPA buffer with added protease inhibitor (ThermoFisher Scientific) and phosphatase inhibitor (PhosSTOP, Roche), and the concentration of the soluble protein fraction was estimated using a Pierce BCA protein assay kit (ThermoFisher Scientific). Protein lysates (15-20μg) were separated by a 4-15% gradient SDS-PAGE gel system (Biorad) and transferred onto a PVDF membrane (ThermoFisher Scientific). Membranes were blocked in 5% non-fat milk in Phosphate-Buffered Saline (PBST) or BSA in Tris-buffered Saline with 0.1% Tween20 (TBST) for 1 hour at room temperature and incubated with primary antibodies at 4 degrees C overnight. After washing with TBST three times, membranes were exposed to appropriate secondary antibodies (anti-rabbit, anti-mouse, and anti-rat) coupled to HRP (Dako) for 1 hour at room temperature. After another set of washes, immunoreactive bands were visualized with the Pierce ECL reagents (ThermoFisher Scientific) and Biorad ChemiDoc XRS+ and ImageLab 6.0.1 software.

### Seahorse assay

In transfected PAECs (20,000 cells/well), oxygen consumption rate and extracellular acidification rate were measured by XFe96 Extracellular Flux Analyzer (Agilent) following exposure to oligomycin (1μM), the uncoupler FCCP (0.5μM), and the Complex I and Complex III inhibitors rotenone (2μM) and antimycin (0.5μM). Basal DMEM contained either high (25mM) or low (1g/L) glucose. Etomoxir (40 µM), a CPT1α (carnitine palmitoyl transferase 1α) inhibitor, was added 15 minutes prior to the assay in a separate experiment to block fatty acid uptake by the mitochondria. Measurements were normalized to protein concentration.

### Hydrogen peroxide assays

Following transfection, cells were counted and re-distributed in a 96-well plate (100,000 cells/well). Intracellular H_2_O_2_ was assessed by Amplex Red Hydrogen Peroxide/Peroxidase Assay kit (Invitrogen). Absorbance (560nm) and/or fluorescence (530nm/590nm) were measured by spectrophotometer (Synergy HTX multimode reader, Biotek). Separately, cultured PAECs were incubated with 2’,7’–dichlorofluorescin diacetate (DCFDA; 1μM) for 10 minutes at 37°C and then assessed by flow cytometry (BD LSRFortessa).

### Endothelin-1 ELISA

Secreted endothelin levels in concentrated serum-free media were assessed by colorimetric change was measured by spectrophotometer (Biotek) using the Endothelin-1 ELISA kit (Enzo Life Sciences).

### Contraction assay

Collagen I (3mg/ml) matrix gel was prepared by mixing collagen I solution (Corning) with a 1/8 volume of 0.1M NaOH and a 1/8 volume of 10X PBS on ice followed by pH adjustment to 7.5 using 0.1M HCL. PASMCs (50,000/well) were embedded in 100μl of matrix gel in a 96-well plate. After 1 hour at 37°C, these matrices were overlaid with 100μl of conditioned PAEC serum-free medium (transfected with siRNA). The endothelin receptor antagonist, ambrisentan (10μM) was supplemented in conditioned media. Media for all groups were changed every 12 hours, and pictures were taken after 2 days. Image J software (NIH) was used to analyze percent contraction ((well diameter – gel diameter)/well diameter).

### Scratch assay

After scratching confluent PAECs with a sterile pipet tip, brightfield images of scratch width were taken at time 0 and 12 hours using an EVOS XL CORE imaging system (Life Technologies). Followed by quantification with NIH ImageJ software (http://rsb.info.nih.gov/ij/) while blinded.

### Tube formation assay

Using the in vitro angiogenesis assay kit (Cultrex), transfected PAECs (30,000 cells/well) were resuspended in 100 µl of basal media and plated in the Matrigel-coated 96-well plate. After 6 hours, tubular structures were photographed using EVOS XL CORE imaging system (Life Technologies) with a 10× magnification. The number of branch points and total tube length were automatically quantified using NIH ImageJ software with Angiogenesis Analyzer plugin (http://rsb.info.nih.gov/ij/).

### Statistics

All data represent three independent experiments and are presented as mean +/-SD unless otherwise specified. Paired samples were compared by a two-tailed Student’s *t*-test for normally distributed data and Mann-Whitney U non-parametric testing for non-normally distributed data. For comparison among greater than two groups, one-way or two-way ANOVA with post-hoc Tukey’s analysis to adjust for multiple comparisons was performed. Significance was defined by a p-value less than 0.05.

## Results

### Acquired FXN deficiency attenuates Fe-S-dependent mitochondrial metabolism

To directly assess mitochondrial respiration, Seahorse extracellular flux analysis was utilized to measure glycolysis (extracellular acidification rate (ECAR)) and oxidative phosphorylation (a component of whole-cell oxygen consumption rate (OCR)) in FXN-deficient PAECs (**Supplemental Figure 1A**). At baseline and following termination of oxidative phosphorylation by the ATP synthase inhibitor oligomycin (indicative of glycolytic capacity), FXN knockdown increased ECAR (**Figure 1A-C**) and expression of glycolytic markers (**Supplemental Figure 1B**) but did not alter OCR (**Figure 1D-E**). Unlike endothelial deficiencies of other Fe-S biogenesis genes such as BOLA3 [10], sustained OCR was not dependent upon fatty acid oxidation, as evidenced by a negligible change in baseline OCR with CPT1α inhibition by etomoxir (**Supplemental Figure 1C**). Instead, by reducing the available glucose, and thus flux through the mitochondria, we found that FXN deficiency decreased baseline and ATP-linked respiration (**Figure 1F-H**), consistent with the notion that oxidative phosphorylation is attenuated due to loss of Fe-S centers in the electron transport chain. Measures of maximal respiratory capacity, proton leak, and non-mitochondrial O_2_ consumption were not significantly different between groups (**Supplemental Figure 1D** and **E**).

**Figure 1.**
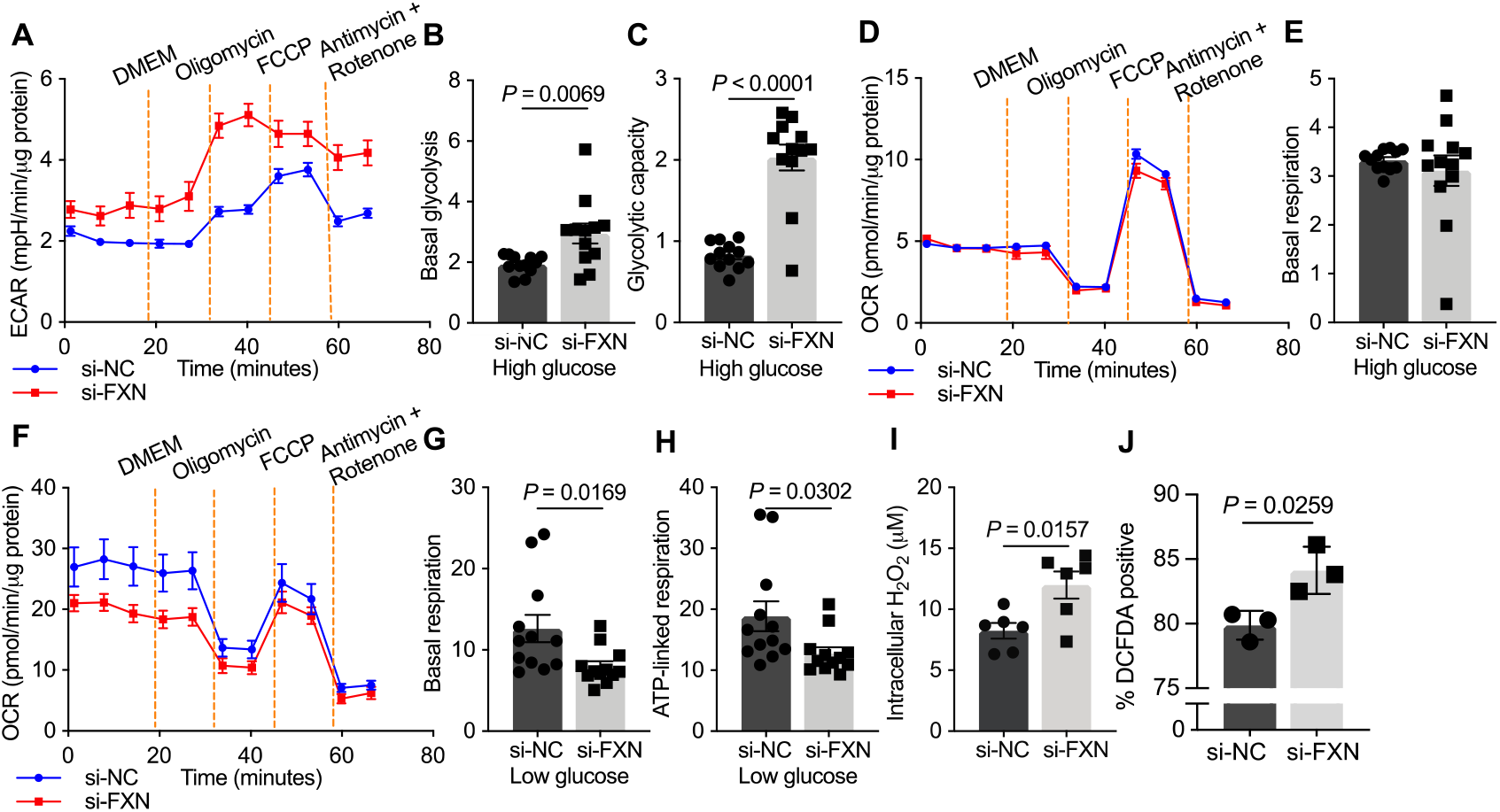
FXN deficiency abrogates mitochondrial respiration and increases oxidative stress. (**A-H**) Seahorse extracellular flux analysis measurements of PAECs transfected with FXN siRNA (red) or negative control (blue) (n=12/group) in response to control (DMEM), the ATP synthase inhibitor oligomycin (1μM), the uncoupler FCCP (0.5μM), and the Complex I and Complex III inhibitors rotenone (2μM) and antimycin (0.5μM). Error bars reflect mean +/-SEM. (**A**) Extracellular acidification rate (ECAR) of PAECs cultured in high glucose (25mM). (**B**) Basal glycolysis (post DMEM). (**C**) Glycolytic capacity (post oligomycin). (**D**) Oxygen consumption rate (OCR) of PAECs in high glucose. Error bars reflect mean +/-SEM. (**E**) Basal respiration (post DMEM). (**F**) OCR of PAECs in low glucose (1g/L). (**G**) Basal respiration (post DMEM). (**H**) ATP-linked respiration (post oligomycin). (**I**) Amplex red colorimetric assay measuring intracellular hydrogen peroxide (H_2_O_2_) in PAECs transfected with FXN siRNA compared to control (n=6/group). (**J**) Flow cytometric percentage of cellular DCFDA fluorescence positivity (n=3/group). Experiments performed at least three separate times and analyzed by two-tailed Student’s *t*-test.

Unlike BOLA3, FXN did not control Fe-S-dependent synthesis of lipoic acid and thus did not affect the glycine cleavage system (**Supplemental Figure 1F**). Corresponding with this metabolic rewiring, FXN deficiency in PAECs increased reactive oxygen species (ROS), as measured by intracellular hydrogen peroxide levels (**Figure 1I-J**). Thus, knockdown of FXN promotes metabolic rewiring and an imbalance in mitochondrial ROS in PAECs. To determine whether mitochondrial oxidative stress directly promotes DNA damage previously demonstrated in endothelial cells with FXN deficiency [17], we employed MnTBAP, a cell-permeable mimetic of superoxide dismutase and peroxynitrite scavenger, to reduce mitochondria-derived ROS. Specifically, using an Amplex red assay, MnTBAP treatment prevented the increase in hydrogen peroxide dependent upon FXN knockdown (**Supplemental Figure 2A**). Next, markers of replication stress, the DNA damage response (DDR), and growth arrest were then measured by immunoblot under similar treatment conditions. Notably, FXN-deficient elevation of these genotoxic stress markers was not prevented by abrogation of mitochondrial ROS production (**Supplemental Figure 2B**), supporting the notion that FXN-driven mitochondrial and nuclear dysfunction occur simultaneously but separately.

### FXN deficiency induces pulmonary artery endothelial dysfunction consistent with PH

Accompanying alterations in mitochondrial metabolism, we found that FXN deficiency promotes endothelial-specific pathophenotypes consistent with PH in addition to DDR-dependent senescence previously described [17]. Namely, in PAECs both transcript and secreted protein levels of the vasoconstrictive mediator endothelin-1 (*EDN1*/ET-1) were increased with FXN knockdown (**Figure 2A-B**). Moreover, the combination of hypoxia and FXN knockdown further elevated this pathologic response (**Supplemental Figure 3A-B**). Demonstrating the role of oxidative stress in this response, EDN1 up-regulation was abrogated by MnTBAP treatment (**Figure 2C**). When cultured in matrigel, primary pulmonary artery smooth muscle cells (PASMCs) exhibited increased contraction after exposure to media from FXN-deficient hypoxic PAECs, which could be reversed with the ET-1 receptor antagonist ambrisentan (**Supplemental Figure 3C**).

**Figure 2.**
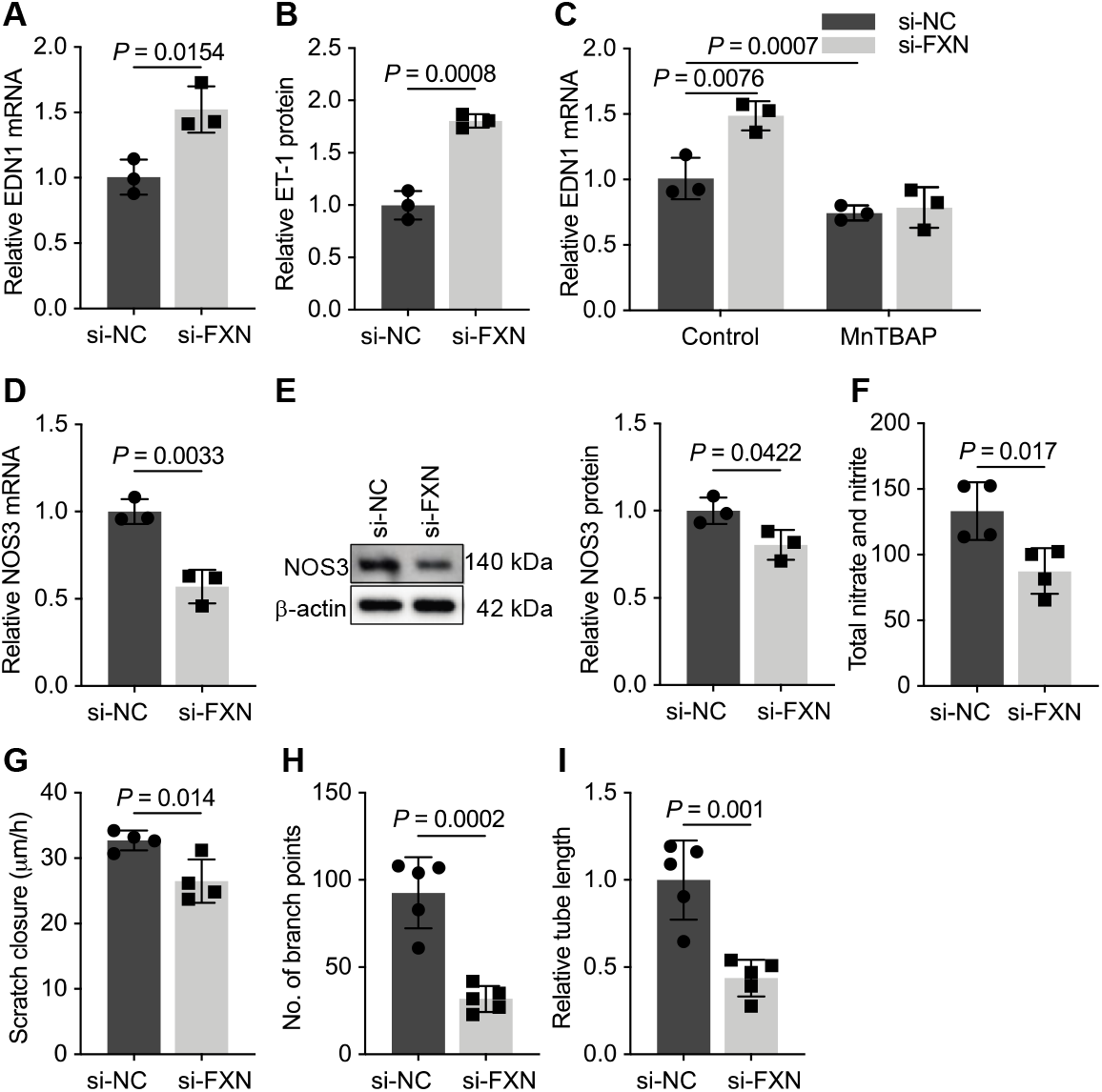
Reduced FXN leads to an imbalance in vasomotor tone effectors and inhibits angiogenesis. (**A** and **B**) RT-qPCR and ELISA measuring endothelin-1 (*EDN1*/ET-1) expression in PAECs transfected with FXN siRNA or negative control (n=3/group). (**C**) EDN1 mRNA levels in FXN-depleted or FXN-replete PAECs treated with the mitochondrial-specific superoxide dismutase MnTBAP (50μM) or vehicle control (n=3/group). (**D** and **E**) RT-qPCR and immunoblot of endothelial nitric oxide synthase (NOS3) expression (n=3/group). (**F**) Total nitrate and nitrite levels (ng/ng) as measured by Griess reagent colorimetric assay (n=4/group). (**G**) Migration rate over 12 hours of FXN-deficient and control PAECs quantified from a scratch assay (n=3/group). (**H** and **I**) Number of branch points and relative tube length reflecting angiogenesis of PAECs cultured in Matrigel (n=4-5/group). Two-tailed Student’s *t*-test with error bars that reflect mean +/-SD. Experiments performed at least three separate times.

FXN knockdown also reduced mRNA and protein expression of nitric oxide synthase 3 (NOS3), the enzyme required for production of the vasodilator nitric oxide (NO) (**Figure 2D-E**). Like EDN1, NOS3 down-regulation was more pronounced during FXN knockdown in combination with chronic hypoxia (**Supplemental Figure 3D-E**). In hypoxia, forced expression of FXN (**Supplemental Figure 3F**) increased NOS3 (**Supplemental Figure 3G**), demonstrating that FXN is both necessary and sufficient for NOS3 expression. Importantly, total nitrite and nitrate levels, a surrogate measurement for NO, were diminished (**Figure 2F**), indicating a decrease of NO production by FXN-deficient PAECs and thus enhancing the vasoconstrictive phenotype. Finally, FXN knockdown, in normoxic and hypoxic conditions, decreased endothelial cell migration during a scratch assay (**Figure 2G, Supplemental Figure 3H**) as well as decreased branch points (**Figure 2H, Supplemental Figure 3I**) and total tube length (**Figure 2I, Supplemental Figure 3J**), consistent with reduced angiogenic potential. In sum, FXN deficiency in PAECs disrupts Fe-S cluster biogenesis, ultimately leading to a dysfunctional endothelium characterized by metabolic stress, an imbalance in vasomotor tone mediators, and diminished angiogenic potential.

### Genetic FXN deficiency mirrors the metabolic dysfunction that drives endothelial dysfunction

To determine whether genetic FXN deficiency produces similar endothelial metabolic dysfunction, inducible pluripotent stem cells (iPSCs) from both male and female patients with FRDA mutations were differentiated into endothelial cells (iPSC-ECs) [20] and compared to gender-and age-matched controls without FXN mutations. FRDA iPSC-ECs exhibited markedly reduced FXN levels (**Supplemental Figure 4A**) and comparable endothelial cell expression markers (**Supplemental Figure 4B-C**). Similar to acquired FXN deficiency, iPSC-ECs with FXN mutations showed an increase in intracellular hydrogen peroxide (**Figure 3A, Supplemental Figure 4D**). This FXN-driven oxidative stress is accompanied by elevated endothelin-1 transcript and secreted protein levels (**Figure 3B-C, Supplemental Figure 4E-F**) as well as reduced NOS3 transcript levels (**Figure 3D, Supplemental Figure 4G**), signifying pathogenic alteration in vasomotor tone effectors similar to those in FXN-deficient primary PAECs.

**Figure 3.**
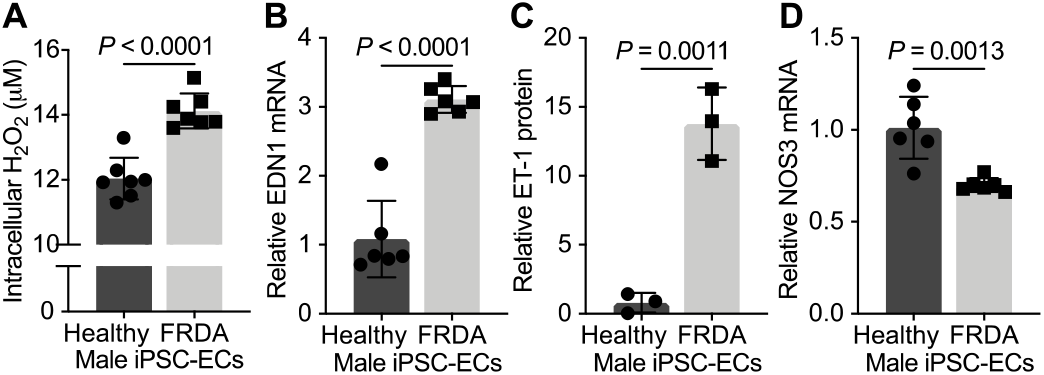
iPS-derived endothelial cells with FXN mutations exhibit similar metabolic and vasomotor pathophenotypes. (**A**-**D**) All phenotypic experiments performed in male age-matched inducible pluripotent stem cell-derived endothelial cells (iPSC-ECs) from a patient with FXN mutations (FRDA) compared to control. (**A**) Amplex red colorimetric assay measuring intracellular hydrogen peroxide (H_2_O_2_) (n=6/group). (**B** and **C**) Relative endothelin-1 transcript (EDN1, n=6/group) and secreted protein (ET-1, n=3/group) expression. (**D**) RT-qPCR of nitric oxide synthase (NOS3) transcript in mutated versus healthy endothelial cells (n=6/group). Two-tailed Student’s *t*-test with error bars that reflect mean +/-SD.

## Discussion

Endothelial mitochondrial dysfunction [1] and DNA damage [21-25] have been separately linked to pulmonary hypertension (PH), but any shared mechanistic regulation of these dynamic endothelial phenotypes in PH was previously undefined. Building upon data that links FXN reduction to Fe-S-specific nuclear damage [17], FXN deficiency disrupted mitochondrial function by preventing Fe-S-dependent glucose oxidation which led to upregulated glycolytic activity and reactive oxygen species. Similar to other Fe-S biogenesis genes [8-10], this mitochondrial oxidative stress resulted in increased EDN1 and decreased NOS3 expression as well as reduced migratory and angiogenic capacity. Our data now cumulatively support that FXN controls not only genotoxic but also metabolic stress resulting in pathophenotypes beyond senescent cell fate (**Graphical Abstract**). By identifying these parallel mechanisms dependent upon FXN, existing vasodilatory or emerging metabolic therapies represent possible tools to help treat PH due to FXN deficiency.

While the shift from oxidative phosphorylation to aerobic glycolysis has been observed in patient and model tissues with FRDA [16], to our knowledge, our data now demonstrate consistent changes in the endothelial cell (**Figure 1**). Whether additional metabolic pathways (*e*.*g*., fatty acid synthesis versus oxidation or the pentose phosphate pathway (PPP)) are dysregulated in FXN-deficient endothelial cells is not yet known. While these pathways do not include Fe-S-containing proteins, loss of appropriate Fe-S-dependent glucose oxidation alone could lead to compensatory alterations in bioenergetic production. For example, the PPP, which yields reductive NADPH and ribose-5-phosphate for nucleotide synthesis, is often augmented in parallel with glycolysis. PPP flux is upregulated in pulmonary vascular cells in multiple PH models [4, 26-28]. Moreover, whether FXN deficiency disrupts mitochondrial biogenesis, dynamics, or even mtDNA integrity requires further investigation.

Our data establish that mitochondria-derived oxidative species do not fully account for the genotoxic stress observed at the replication fork (**Supplemental Figure 2B**). In tandem with our data on disruption of the Fe-S-containing polymerases in FXN-deficient endothelial cells, these data support the notion that there may exist two separate organelle-specific processes dependent upon Fe-S cluster loss. However, this data does not preclude the possibility of bi-directional signaling between organelles, particularly over the course of disease. In particular, FXN-dependent metabolic changes may contribute to the senescent cell fate previously reported in the FXN-deficient endothelium [17]. Mitochondrial dysfunction-associated senescence (MiDAS) represents another stress response pathway like telomere instability or the DNA damage response that converges on permanent growth arrest, but the mechanisms remain incompletely defined [29]. We acknowledge that further experimentation is needed to determine the importance of mitochondrial versus nuclear stress in PH, as these may provide insight into the value of metabolic versus genomic treatment intervention. Moreover, the potential reciprocal relationships between these organelles may favor targeting FXN directly and require the identification of new drugs in addition to potentially emerging epigenetic therapies for FRDA (*e*.*g*., HDAC inhibitors [30-32]).

While our data mirror the same endothelial pathophenotypes demonstrated with other Fe-S biogenesis gene deficiencies with Warburg-like metabolic shifts [8-10], only FXN-dependent endothelin-1 up-regulation could be reversed by targeting mitochondrial superoxide (**Figure 2C**). Therefore, oxidative stress may not account for all of the Fe-S-dependent phenotypes presented here; the precise mechanisms that result in changes in vasomotor tone and angiogenesis remain incompletely defined. In the context of FXN-dependent down-regulation of NOS3 expression (**Figure 2D-E**), a possible explanation could center on FXN-specific inhibition of ferrochelatase activity [33], the Fe-S-dependent, rate-limiting enzyme in heme synthesis; thus FXN deficiency may impair heme production and ultimately reduce the expression and activity of NOS3, a heme-containing enzyme [34], resulting in diminished nitric oxide (NO) production. Separately, because nitric oxide can directly bind and damage Fe-S clusters, endothelial cell NOS3 may be inhibited by another undefined feedback mechanism in conditions of FXN deficiency and reduced Fe-S biogenesis [35].

Regardless, Fe-S-mediated changes in effectors of vasomotor tone suggest current PH-specific vasodilatory therapies that inhibit ET-1 and enhance NO signaling may be effective in circumstances of endothelial FXN deficiency, particularly in FRDA. However, certain types of PH patients do not respond to vasodilatory therapy, including most Group 2 and Group 3 PH patients; yet, FXN deficiency has been identified across Group 1-3 PH subtypes. This may be reflective of the fact that FXN-deficient endothelial cells only represent a subpopulation rather than a predominance of the endothelium in the lung vasculature. In this way, their senescence-dependent inflammatory phenotype may be more important in the pathology of PH than their contribution to vasoconstrictive signaling. Of note, reduced NO availability has been appreciated in senescent endothelial cells [36] while repletion of NO signaling prevented senescence [37]. While the onset of FXN-dependent alterations in vasomotor tone effectors predate the onset of senescence, the vasodilatory therapies may serve a dual purpose that includes preventing endothelial senescence over the course of the disease.

In conclusion, these findings support FXN as a lynchpin connecting Fe-S-dependent endothelial metabolic reprogramming and genotoxic stress critical to PH development, reinforcing the importance of Fe-S biology in endothelial cell function and offering therapeutic targets in PH.

## Abbreviations

Fe-S: Iron-sulfur
PH: Pulmonary hypertension
FXN: Frataxin
FRDA: Friedreich’s ataxia
ROS: Reactive oxygen species
HIF-α: hypoxia-inducible factor alpha
iPSC-EC: inducible pluripotent stem cell-derived endothelial cell
PAEC: pulmonary artery endothelial cell
PASMC: pulmonary artery smooth muscle cells
ECAR: Extracellular acidification rate
OCR: Oxygen consumption rate
EDN1/ET-1: Endothelin-1,
NOS3: Nitric oxide synthase 3,
NO: Nitric oxide
PPP: Pentose phosphate pathway

## Acknowledgments

Portions of this manuscript and data were included in M.K.C.’s unpublished doctoral dissertation (Culley, Miranda (2020) *Frataxin deficiency coordinates iron-sulfur dependent metabolic and genomic stress to promote endothelial senescence in pulmonary hypertension*. Doctoral Dissertation, University of Pittsburgh). This study used inducible pluripotent stem cell samples from patients with FRDA (GM23404, GM23913) in the NIGMS Human Genetic Cell Repository at the Coriell Institute for Medical Research. We also thank Y. Lu, S. Annis, and M. Reynolds for their technical assistance.

## Author Contributions

M.K.C.: Conceptualization, Investigation, Formal Analysis, Visualization, Writing - Original Draft; M.M.: Writing - Original Draft; J.Z.: Investigation, Validation, Formal Analysis, Writing - Review & Editing; D.P.: Investigation, Validation, Writing - Review & Editing; Y.Y.T.: Resources, Writing - Review & Editing; Y.T.: Resources, Writing - Review & Editing; S.S.: Methodology, Resources, Writing - Review & Editing; M.G.: Methodology, Resources, Writing - Review & Editing; M.R.: Methodology, Resources, Writing - Review & Editing; T.B.: Methodology, Writing - Review & Editing; and S.Y.C.: Conceptualization, Supervision, Project administration, Funding acquisition, Writing - Original Draft.

## Financial support

This work was supported by National Institutes of Health grants F30 HL139017 (M.K.C.), 5R00 HL135258 (M.G.), 5U01 HL10739302 (M.R.), R01 HL124021 and HL122596 (S.Y.C). Additional support includes the Friedreich’s Ataxia Research Alliance general research grant and American Heart Association grant 18EIA33900027 (S.Y.C.).

## Disclosures

S.Y.C. has served as a consultant for Acceleron Pharma and United Therapeutics. S.Y.C. has held research grants from Actelion, Bayer, and Pfizer. S.Y.C. is a director, officer, and shareholder of Synhale Therapeutics. S.Y.C. and T.B. hold patents and has filed patent applications regarding the therapeutic targeting of metabolism and senescence in pulmonary hypertension.

