## Supplemental Data for "Frataxin deficiency disrupts mitochondrial respiration and pulmonary endothelial cell function"

### Supplemental Figures

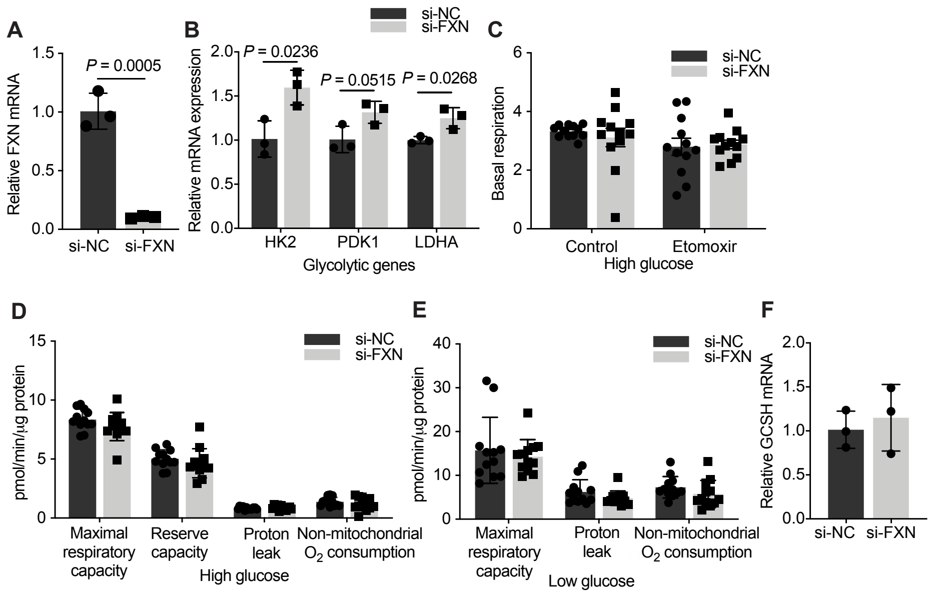


**Supplemental Figure 1**

**Supplemental Figure 1. Effects of FXN deficiency on endothelial metabolism.** (**A**) FXN transcript expression by RT-qPCR following siRNA transfection in PAECs (n=3/group). (**B**) RT-qPCR expression analysis of glycolytic markers: hexokinase 2 (HK2), pyruvate dehydrogenase kinase 1 (PDK1), and lactate dehydrogenase A (LDHA) (n=3). (**C**) Basal respiration with or without etomoxir (40μM) in high glucose media (n=12/group). Error bars reflect +/- SEM. (**D**) Oxygen consumption rate (OCR; pmol/min/μg protein) measurements of maximal respiratory capacity, reserve capacity, proton leak, and non-mitochondrial O2 consumption by Seahorse assay in high glucose media (25mM) (n=12/group). Error bars reflect +/- SEM. (**E**) OCR measurements of maximal respiratory capacity, proton leak, and non-mitochondrial O2 consumption by Seahorse assay in low glucose media (1g/L) (n=12). Error bars reflect +/- SEM. (**F**) GCSH mRNA levels in FXN-deficient PAECs compared to negative control (n=3). Two-tailed Student's *t*-tests were performed, and error bars reflect +/- SD unless otherwise specified.

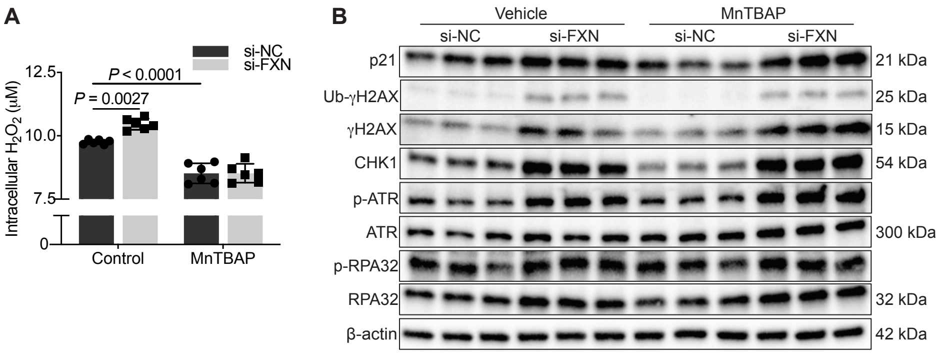


**Supplemental Figure 2**

**Supplemental Figure 2. FXN-dependent mitochondrial oxidative stress does not induce nuclear genotoxic stress.** (**A**) Intracellular $H_2O_2$ quantified by Amplex red assay following treatment with the mitochondrial-specific superoxide dismutase MnTBAP (50μM, ≥24 hours) in PAECs with and without silencing of FXN (n=6/group). Data presented as mean +/- SD and analyzed by two-way ANOVA (Tukey's post hoc analysis). (**B**) Representative immunoblot of replication stress (p-RPA32/RPA32), DNA damage response (p-ATR/ATR, CHK1, Ub-γH2AX/γH2AX), and growth arrest markers ($p21^{Cip}$) in FXN-deficient PAECs in response MnTBAP (n=3/group).

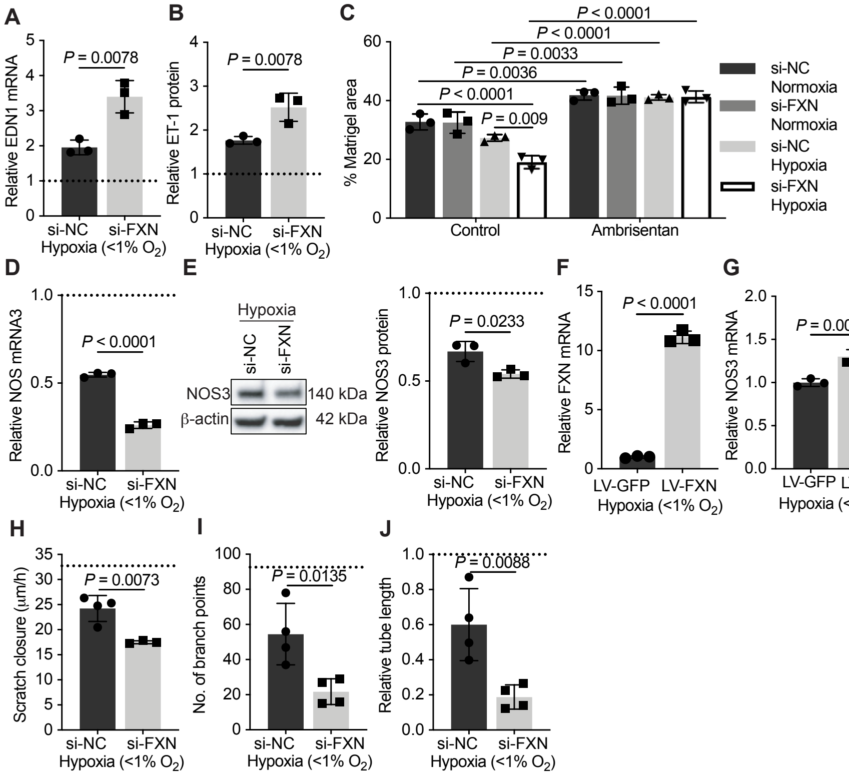


**Supplemental Figure 3**

**Supplemental Figure 3. Chronic hypoxia worsens FXN-dependent endothelial dysfunction.** (**A**-**H**) Experiments reflect PAECs transfected with FXN siRNA compared to control (si-NC) and exposed to hypoxia (≥24 hours) unless otherwise specified. The dotted line represents mean data in PAECs without FXN silencing in normoxic conditions (21% $O_2$). (**A** and **B**) Relative endothelin-1 mRNA (EDN1) and secreted protein (ET-1) expression (n=3/group). (**C**) Contraction of human pulmonary artery smooth muscle cells (PASMCs) seeded in a collagen matrigel measured as a percentage of well area compared to baseline area following treatment with conditioned media from PAECs with and without FXN knockdown, exposed to hypoxic or normoxic conditions, and treated with the endothelin receptor antagonist, ambrisentan (≥ 48 hours, 10μM) (n=3/group). (**D** and **E**) RT-qPCR and immunoblot of relative nitric oxide synthase (NOS3) levels (n=3/group). (**F**) RT-qPCR of FXN transcript following transduction with lentiviral vectors containing FXN (LV-FXN) versus GFP control (LV-GFP) (n=3/group). (**G**) Rescue of NOS3 mRNA expression in hypoxic PAECs with forced FXN expression (n=3/group). (**H**) Rate of cellular migration by measurement of scratch closure (n=3-4/group). (**I** and **J**) Tube formation assay measuring number of branch points and tube length reflecting angiogenesis (n=4/group). Two-tailed Student's *t*-test or two-way ANOVA (Tukey's post hoc analysis) with error bars that reflect mean +/- SD. Experiments repeated three times.

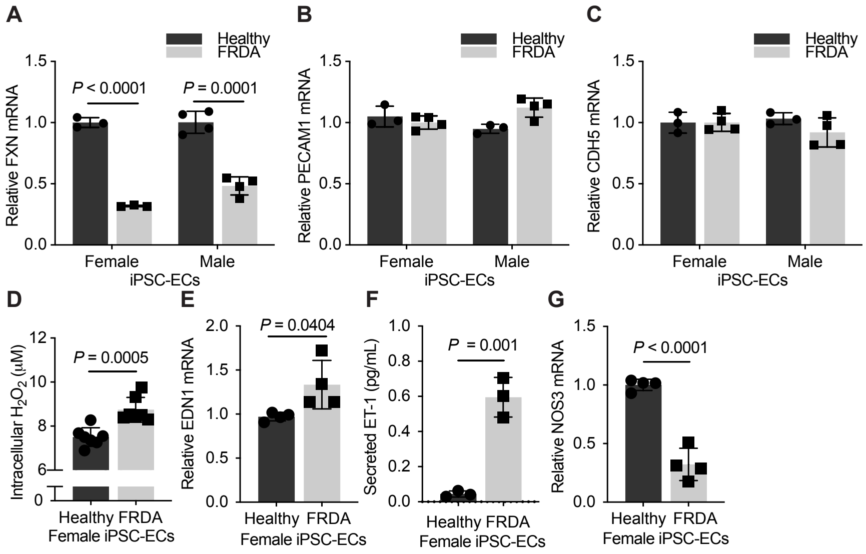


**Supplemental Figure 4**

**Supplemental Figure 4. Characterization of inducible pluripotent stem cell-derived endothelial cells from patients with Friedreich's ataxia.** (**A**) RT-qPCR of FXN levels in iPSC-ECs from female and male patients with Friedreich's ataxia (FRDA) compared to healthy iPSC-ECs (n=3-4/group). (**B** and **C**) Transcript expression of endothelial-specific markers PECAM1 and CDH5 (n=3-4/group). (D-F) Phenotypic experiments performed in female age-matched iPSC-ECs. (**D**) Amplex red assay measuring intracellular $H_2O_2$. (**E** and **F**) Relative endothelin-1 transcript (EDN1, n=6/group) and secreted protein (ET-1, n=3/group) expression. (**G**) RT-qPCR of nitric oxide synthase (NOS3) transcript in mutated versus healthy endothelial cells (n=4/group). Two-tailed Student's *t*-test with error bars that reflect mean +/- SD.
